# *Enterococcus faecium* genome dynamics during long-term asymptomatic patient gut colonization

**DOI:** 10.1101/550244

**Authors:** Jumamurat R. Bayjanov, Jery Baan, Malbert R.C. Rogers, Annet Troelstra, Rob J.L. Willems, Willem van Schaik

## Abstract

**Background:** *E. faecium* is a gut commensal of humans and animals. In addition, it has recently emerged as an important nosocomial pathogen through the acquisition of genetic elements that confer resistance to antibiotics and virulence. We performed a whole-genome sequencing based study on 96 multidrug-resistant *E. faecium* strains that asymptomatically colonized five patients with the aim to describe the genome dynamics of this species.

**Results:** The patients were hospitalized on multiple occasions and isolates were collected over periods ranging from 15 months to 6.5 years. Ninety-five of the sequenced isolates belonged to *E. faecium* clade A1, which was previously determined to be responsible for the vast majority of clinical infections. The clade A1 strains clustered into six clonal groups of highly similar isolates, three of which entirely consisted of isolates from a single patient. We also found evidence of concurrent colonization of patients by multiple distinct lineages and transfer of strains between patients during hospitalisation. We estimated the evolutionary rate of two clonal groups that colonized a single patient at 12.6 and 25.2 single nucleotide polymorphisms (SNPs)/genome/year. A detailed analysis of the accessory genome of one of the clonal groups revealed considerable variation due to gene gain and loss events, including the chromosomal acquisition of a 37 kbp prophage and the loss of an element containing carbohydrate metabolism-related genes. We determined the presence and location of twelve different Insertion Sequence (IS) elements, with IS*Efa5* showing a unique pattern of location in 24 of the 25 isolates, suggesting widespread IS*Efa5* excision and insertion into the genome during gut colonization.

**Conclusions:** Our findings show that the *E. faecium* genome is highly dynamic during asymptomatic colonization of the patient gut. We observe considerable genomic flexibility due to frequent horizontal gene transfer and recombination, which can contribute to the generation of genetic diversity within the species and, ultimately, can contribute to its success as a nosocomial pathogen.

## Background

In recent decades, *Enterococcus faecium* has emerged as an important multidrug-resistant nosocomial pathogen. It is a major cause of hospital-acquired infections such as bacteraemia, urinary tract infection and endocarditis [1–4]. Furthermore, enterococcal infections contribute to patient mortality, increased length of hospital stay of patients and higher healthcare costs [5]. Infections caused by *E. faecium* are difficult to treat due to the large repertoire of acquired antibiotic resistance determinants, of which vancomycin resistance is arguably the most problematic [6, 7].

The species *E. faecium* consists of distinct subpopulations or ‘clades’ [8–10]. A deep phylogenetic split distinguishes clades A and B from each other [11], with clade B containing most human commensal isolates. Clade A was further sub-divided in clade A1 and A2 [8]. Clade A1 contains the vast majority of strains isolated from clinical settings, and overlaps with the previously identified *E. faecium* sub-population Clonal Complex 17 [9, 12]. The polyphyletic clade A2 is enriched for strains that were isolated from domestic animals and livestock [8, 10]. While vancomycin resistance can be found among strains from both clade A1 and clade A2, clade A1 strains are almost always resistant to ampicillin, while strains from other clades are mostly ampicillin-susceptible [12].

*E. faecium* is a genetically dynamic organism with an open pan-genome [8, 13, 14]. Genomic changes in *E. faecium* are mostly driven by recombination and horizontal gene transfer (HGT), rather than by mutation [15]. Due to frequent HGT, *E. faecium* strains that have highly similar core genomes can have substantial differences in their accessory genomes [8, 14, 16]. Insertion Sequence (IS) elements are abundant in *E. faecium* genome [16]. IS elements are short transposable segments of DNA that can have an important role in shaping a bacterial genome. Insertion events can lead to disruption of promoters, coding sequences or operon structures. In addition, they can catalyze genomic rearrangements including deletions, inversions, and duplications in bacterial genomes [17]. Complete genome sequences revealed that dozens of IS elements are scattered around the chromosome and plasmids of clinical *E. faecium* isolates [18, 19]. A number of IS elements, most notably IS*16*, are associated with clade A1 strains and have been hypothesized to contribute to the adaptation of *E. faecium* to the hospital environment [8, 16].

Patients that have been hospitalized for prolonged periods of time are potential reservoirs for drug-resistant *E. faecium* strains. Generally, infection by *E. faecium* is preceded by asymptomatic gut colonization by a resistant clade A1 strain [20, 21]. Patients that have been colonized by *E. faecium* can contaminate both their immediate surroundings and healthcare workers, leading to outbreaks [21, 22]. The ability of *E. faecium* to survive on inanimate objects creates an environmental reservoir of multidrug-resistant strains in hospital wards and makes outbreaks with *E. faecium* a challenge to control [23].

Recent studies have used whole-genome sequencing (WGS) to trace transmission events of *E. faecium* between patients in hospital wards and between hospitals [10, 24, 25]. Recently, the relatedness of *E. faecium* strains from bloodstream infections, the gut and the immediate environment of four patients that were hospitalized for up to two months was studied using WGS [21]. Here, we present an analysis of the genome dynamics of vancomycin- and ampicillin-resistant *E. faecium* during asymptomatic gut colonization of five patients for periods ranging from 15 months to 6.5 years. We describe the evolutionary trajectories, including the roles of gene gain and loss events, and IS-element excision and insertion that shape the genome of *E. faecium*.

## Results

### Isolate collection and patient hospital stay

This study used vancomycin- and ampicillin-resistant *E. faecium* (VRE and ARE, respectively) isolates that were collected and stored in the period 2001 - 2008 as part of routine diagnostics and infection prevention interventions at the University Medical Center Utrecht, the Netherlands (figure 1). Analysis of the collected isolates with anonymized patient data, showed that for five patients multidrug-resistant *E. faecium* isolates were collected over a period of >1 year. We sequenced the genomes of 96 isolates, all of which were determined to be ampicillin-resistant using a previously described method [26]. Using Abricate [27], we found that 38 and 21 isolates carried the *vanA* or *vanB* operon. Further information on the antibiotic resistance profiles of the strains sequenced in this study is provided in Supplementary Table 1. The time span between the first and the last isolate collected from a single patient ranges from 15 months (patient B) to 6.5 years (patient C).

**Figure 1:**
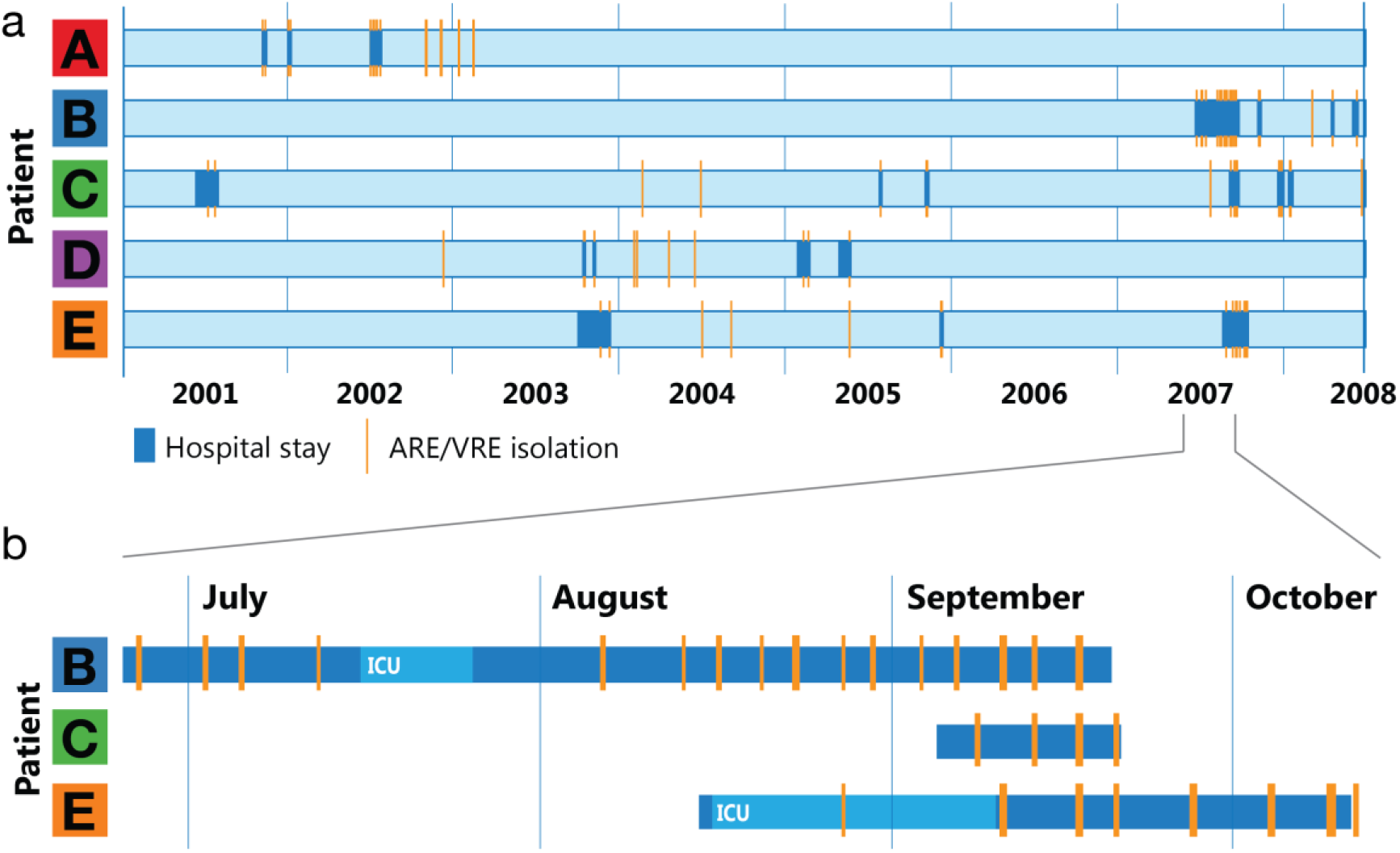
(a) Timeline of hospital stay for 5 patients (A-E) and the time points at which multi-drug-resistant *E. faecium* strains were isolated during routine screening between 2001 and 2008. (b) Detail for patients B, C, and E, showing the overlap in their hospital stay in 2007 and the associated ARE/VRE isolations. Dark blue: patient hospital stay; orange: ARE/VRE-positive screening; ICU: patient in an intensive care unit. If an isolation time point does not overlap with hospital stay, the screening was performed at home as part of outbreak control studies.

### Genetic diversity of *E. faecium* patient isolates

To be able to place the collected isolates in the larger *E. faecium* population, we created a SNP-based, recombination-filtered phylogenetic tree using the 96 genomes sequenced in this study and 70 previously described *E. faecium* genome sequences that represent the global *E. faecium* population [8]. This phylogenetic tree is based on 1448 core genes and a total of 77,909 SNPs. Out of the 96 patient isolates, 95 clustered into clade A1, a clade of hospital-associated *E. faecium* strains (Supplementary figure 1). The remaining isolate clustered with strains that were previously assigned to clade A2.

The relatively large diversity of the 70 publicly available *E. faecium* genome sequences, limited the resolution of the phylogenetic relationships between the patient isolates. We therefore created a second tree, with only the 95 clade A1 patient isolates in this study, supplemented with 19 genome sequences of clade A1 strains that were previously sequenced [8]. This tree was based on 1805 core genes, with 5092 SNPs and allowed us to accurately interpret the similarities between the hospital isolates (figure 2). The phylogenetic tree of the clade A1 strains revealed 6 groups of closely related isolates. Three of these groups (1, 2 and 3) contained only isolates from a single patient (A, B and D, respectively). While additional isolates of patients B and D were present in other groups, patient A isolates clustered exclusively in group 1. In group 4, isolates clustered closely together despite being from three different patients (B, C and E). By analysis of dates and locations of hospitalization of these patients, we found that patients B, C and E were simultaneously present in a hospital ward (figure 1b), suggesting that we captured a small outbreak with this set of isolates. Groups 5 and 6 are two small, highly similar clusters of isolates. The isolates in group 5 originate from 3 patients (C, D, E) which were isolated between October 2003 and February 2004. In group 6, isolates from patients C and E that were isolated between September 2007 and January 2008 cluster together. Both of these groups may thus also reflect transmission of strains between different patients.

**Figure 2:**
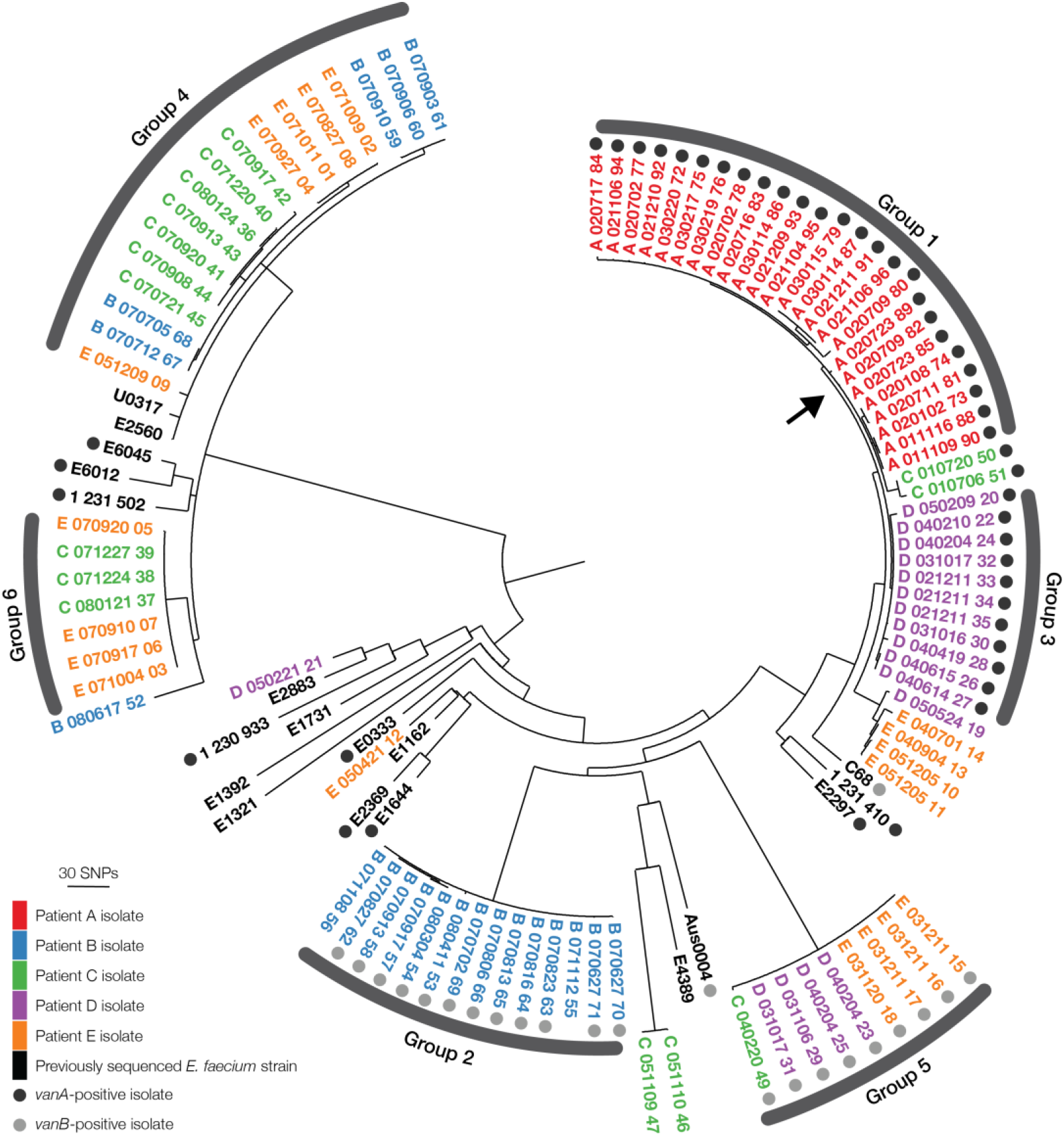
Phylogenetic tree of clade A1 isolates. This maximum-likelihood tree includes 95 of the 96 genome sequences generated in this study and 19 publicly available *E. faecium* genome sequences. The core genome alignment consisted of 2,295,725 nucleotides. The position of strain A_020709_82 is marked with an arrow. The genome of this isolate was sequenced and assembled to completion using a combination of short- and long-read sequencing for use as a reference genome in further analyses. Genome sequences are coded as follows: the letter represents the patient, the six number code represents the data of isolation in year-month-day format, the final number is the unique identifier for each genome sequence. Colours indicate the patient this isolate was taken from. Black and gray marks indicate the presence of the *vanA* or *vanB* vancomycin resistance operon in the genome, respectively.

We also found that patients can be colonized by different populations of *E. faecium* at the same time, as shown by the genetic diversity found in strains that were isolated on the same date, e.g. those on 4 February 2004 (D_040204_23, D_040204_24 and D_040204_25).

We determined whether a temporal signal is present in the *E. faecium* genome sequences in each individual group. The temporal signal was defined using Path-O-Gen [28], which plots the time at which each isolate was identified versus the distance to the root of the tree. A temporal signal (R^2^>0.3), was only found in groups 1 and 4. Analysis by BEAST resulted in estimated mutation rates for groups 1 and 4 of 4.2 x 10^−6^ (with 95% highest posterior density (HPD) of [2.2 x 10^−6^, 6.3 x 10^−6^]) and 8.4 x 10^−6^ (with 95% HPD of [4.7 x 10^−6^, 1.2 x 10^−5^]) substitutions per nucleotide per year, being equivalent to 12.6 (6.6 - 19.8) SNPs/genome/year and 25.2 (14.1 - 36.0) SNPs/genome/year, respectively. The lack of temporal signal in groups 2, 3, 5, and 6 is likely caused by the relatively low number of strains in these groups. Because all group 1 isolates originate from the same patient, they provide a unique opportunity to study the genome dynamics of *E. faecium* during long-term asymptomatic patient gut colonization. Hence, we focused further analyses on the strains from this group.

### The accessory genome of group 1 strains

A total of 74 orthologous genes (OGs) were found to be differentially present in the genomes of the group 1 isolates (figure 3a). Hierarchical clustering showed that most of the OGs were part of larger groups of OGs that showed the same presence-absence pattern across the genome sequences, suggesting that they are genetically linked. Further analysis revealed that the clustered OGs were co-located on contigs.

**Figure 3:**
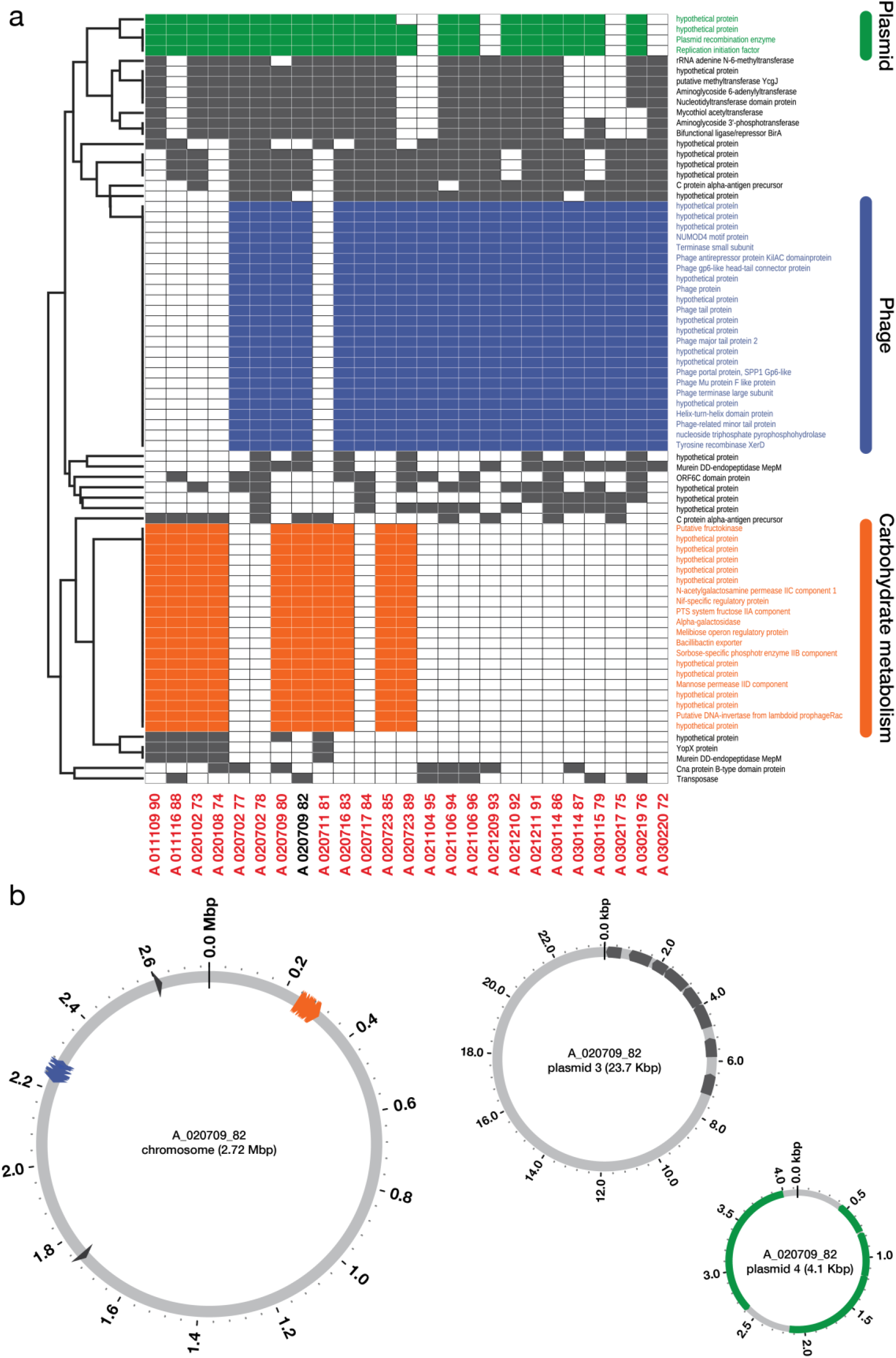
The accessory genome of group 1 isolates. (a) Plot showing the differentially present genes in the different isolates, ordered chronologically. Colours indicate gene clusters that are variably present or absent and are annotated on the basis of their predicted function or origin. (b) The differentially present genes mapped onto the A_020709_82 genome, with colours corresponding to gene clusters in panel a. Chromosome and plasmid sizes not shown to scale.

The two largest variably present clusters are phage-related OGs (cluster 1), and OGs related to carbohydrate metabolism (cluster 2). Cluster 1 contains 24 OGs, of which 10 are annotated as being hypothetical proteins. The annotations of the remaining genes suggested a phage origin of this element as they included tail and terminase protein-encoding genes (figure 3a). Cluster 2 comprised several genes that are related to carbohydrate transport. Neither of the clusters contained genes related to antimicrobial resistance.

When aligning these gene clusters to the original collection of 166 genomes (96 genomes sequenced in this study and 70 genomes representing global *E. faecium* diversity) using BLAST, we find that they are mostly found in the newly sequenced isolates (Supplementary figure 1), with cluster 1 being found in 42 genomes, of which 38 were sequenced as part of this study. Cluster 2 was present in 28 genomes, of which 26 were sequenced here.

To further investigate the genetic linkage of these variably present clusters in the accessory genome of group 1 strains, we fully sequenced the genome of isolate A_020709_82, combining Illumina reads with long reads generated via Oxford Nanopore’s MinION platform to complete the genome assembly. The A_020709_82 strain has most of the genes of the accessory genome that are variably present among group 1 strains, including the two largest groups of OGs. The A_020709_82 strain has a chromosome of 2,740,566 nucleotides and 4 plasmids, ranging in size from 222 kbp to 4 kbp (figure 3b). By mapping all the differentially present OGs onto the A_020709_82 reference genome sequence, we found that the clustered OGs were located in close proximity to one another in the chromosome. A third, smaller variably present cluster consisting of 4 OGs, was found to be representing a 4.1 kbp plasmid that is lost in its entirety in 4 of the isolates, in a presence/absence pattern unrelated to that of the two larger clusters. To assess whether the differences in accessory genome sequences influence the fitness of the 25 group 1 isolates, we determined their *in vitro* maximum growth rates but found no statistically significant differences between the strains with different accessory genomes (Supplementary Figure 2).

### Dynamics of IS elements in a clonal *E. faecium* population

We identified 12 different IS elements in the genome of A_020709_82. To identify the diversity and location of IS elements in the other strains from group 1, we used ISmapper [29] with the A_020709_82 genome as a reference and the sequencing reads of the other genomes in group 1 (figure 4). The positions of two IS elements (IS*16* and IS*6770*) are fixed in all 25 isolates. The IS-element IS*Efa5* exhibited a particularly large diversity, having between 17 and 27 copies per genome. Twenty-four out of the 25 isolates in group 1 have a unique pattern, suggesting frequent excision and integration events of this IS element. The remaining 9 IS elements showed an intermediate amount of diversity.

**Figure 4:**
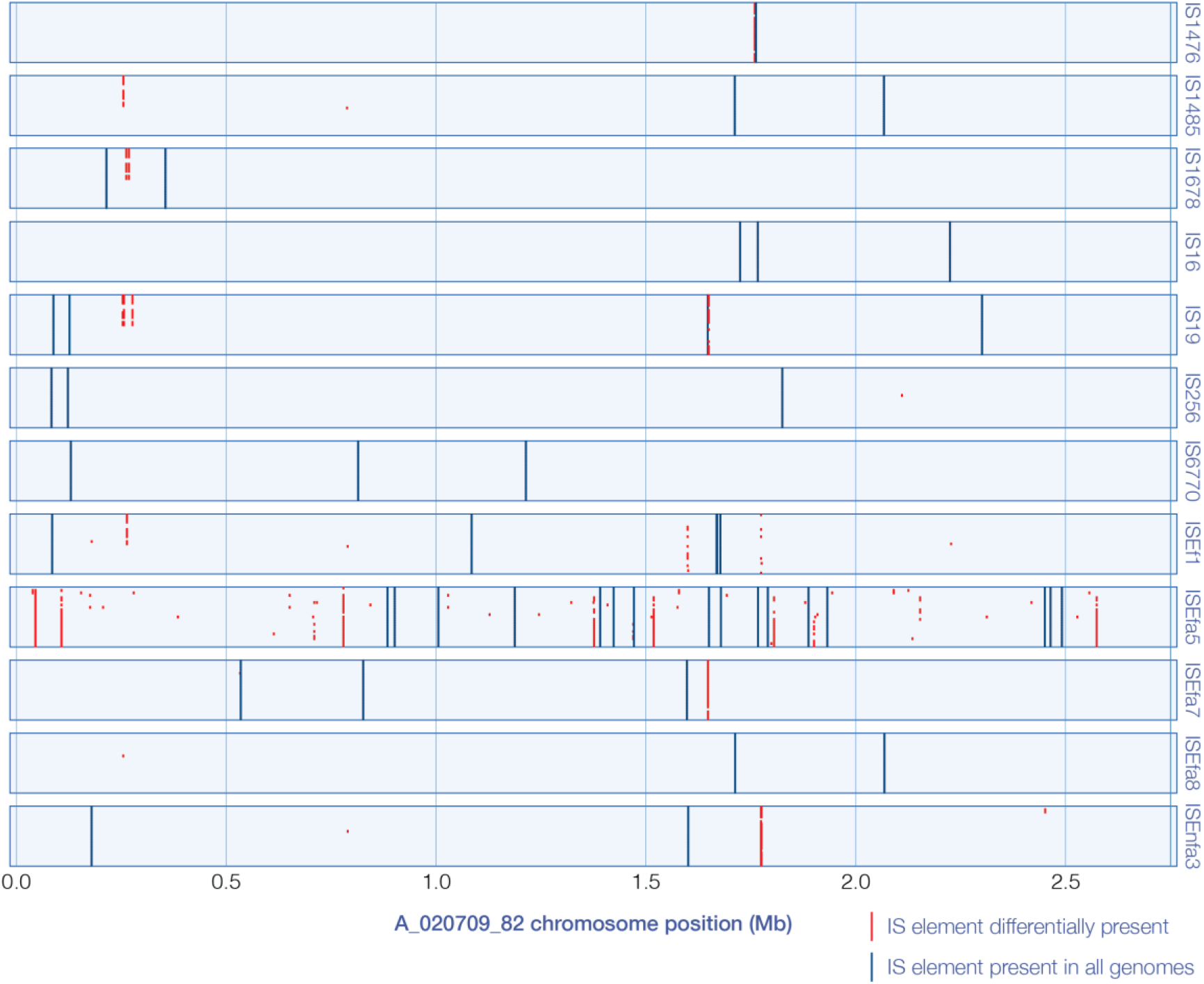
Variable presence of IS elements in a clonal population of *E. faecium* during asymptomatic gut colonization. Overview of the different IS elements found in the genomes of the 25 clonal patient A isolates, plotted on the chromosomal sequence of isolate A_020709_82. A total of 12 different IS elements are found in this group. Blue marks indicate the presence of that IS-element in all isolates. Red marks indicate that the IS element is present in the indicated isolate, but not in all isolates. Each row of an individual IS element represents a single isolate, with the oldest isolate on top and the newest at the bottom.

## Discussion

In this study, we use a collection of *E. faecium* carriage strains that were isolated from patients that were repeatedly admitted to a hospital over a time period ranging from 15 months to 6.5 years. Out of 96 isolates, 95 clustered to the hospital-associated A1 clade, which is expected given their source as ampicillin-resistant clade A1 strains cause the majority of hospital-acquired infections, and are rarely carried by humans in community settings [8, 30]. The patients likely acquired these isolates during their hospital stay and were carriers for extended periods of time. Previous work has shown that ampicillin-resistant *E. faecium* clones can persist in the gut microbiota for several months after discharge from hospital [31], during which time further spread can occur.

The mutation rate we find in both the clonal group 1 and non-clonal group 4 (12.6 and 25.2 substitutions/genome/year, respectively) are in line with previously described values for similar *E. faecium* populations [13, 32]. Others have described rates of up to one order of magnitude higher [8, 21, 25]. This difference is postulated to be caused by increased genetic drift within patients along with a limited time for purifying selection to act on a population, leading to the incomplete removal of strains with mildly deleterious mutations [21, 33]. Our group 1 estimation in particular can be assumed to be a better approximation of the background mutation rate of *E. faecium* given their clonality, the absence of enterococcal disease in the source patient, and the longer time over which they were collected. However, it is also possible that the large differences in the mutation rate of different *E. faecium* clones reported in literature are a true biological signal. As in the Gram-negative gut commensal *E. coli*, *E. faecium* clones with a higher mutation rate may be able to more rapidly adapt to novel environments while negatively impacting their transmissibility and ability to recolonize similar hosts [34].

The pangenome of *E. faecium* has previously been determined to be essentially open, meaning that it can easily acquire novel genes by horizontal gene transfer [8, 14]. This ability to acquire DNA was recently vividly illustrated by the description of a bovine *E. faecium* strain that had acquired a gene cluster encoding a botulinum-like neurotoxin [35]. In group 1 strains we observed a number of gene gain and loss events. The earliest isolates in group 1 carry a gene cluster that is predicted to be involved in carbohydrate metabolism, while later strains lose this element and acquire a phage. Five isolates carry both the phage element and the carbohydrate metabolism gene cluster, which shows that carriage of both elements is not mutually exclusive. While we did not observe differences in the *in vitro* growth rates of strains with different combinations of the carbohydrate metabolism and phage element, their presence might affects the strains’ fitness in colonizing gut of this patient. It is possible that the changes in the accessory genome allow the clone to optimally adapt to colonize in the context of the patient’s gut microbiota.

As described in previous studies, there is an abundance of IS elements in the *E. faecium* genome [16]. We find that some IS elements, such as IS*256*, IS*6770* and the clade A1-associated IS*16* [8], show little to no variation in insertion location and number in the genomes of patient A isolates. IS*16* was previously proposed to confer a degree of genomic flexibility to the hospital-adapted sub-population of *E. faecium* that could contribute to its success as a nosocomial pathogen[16]. However, the fixed position of IS*16* in the group A isolates appears to contradict a major role for this IS element in shaping the *E. faecium* genome. It may be more likely that IS*16* has entered the *E. faecium* population when the hospital-adapted sub-population (clade A1) first emerged and has since been spreading vertically in this population. Conversely, we find a large number of IS*Efa5* copies in the genome of group 1 strains and evidence for frequent excision and insertion events. IS*Efa5* was first described as part of Tn*1546*-like elements, which are responsible for VanA-type vancomycin resistance, in South American *E. faecium* isolates [36, 37], but it was later found in European [38] and Australian strains [39] as well. In the whole genome sequence of strain Aus0085 25 copies of IS*Efa5* were found [39]. Its high copy number in *E. faecium* strains and the evidence provided in this study for frequent integration and excision events, suggests that IS*Efa5* may be contributing significantly to the genomic flexibility of the species.

Our observation that patients can be colonized by multiple strains simultaneously is in line with previous studies [21, 33, 40]. Concurrent colonization by multiple clones can have an important impact on infection prevention efforts if only single colonies are selected for further typing. Potentially pathogenic or multidrug-resistant strains can then be inadvertently missed, leading to the erroneous reconstruction of transmission networks. When isolates are missed, transmission networks may also be reconstructed erroneously [21], making outbreak control more challenging. This is illustrated by the small outbreak we detected in our dataset, where patients B, C and E appear to be colonized by isolates with high inter-patient similarity, as well as more different ones. Sampling and typing of multiple colonies when performing screening for colonization by multi-drug resistant *E. faecium* is thus required to capture the full within-patient diversity of this organism. In addition, the use of metagenomic shotgun sequencing, combined with tools to reconstruct microbial genomes and resolve strains [41] may become a useful alternative to culture-based approaches to determine the presence of different *E. faecium* clones in the gut microbiome.

## Conclusions

Our findings show that the *E. faecium* genome is highly dynamic during asymptomatic colonization of the patient gut. We demonstrate *E. faecium*’s remarkable genomic flexibility, which is characterized by frequent gene gain and gene loss due to horizontal gene transfer and recombination and the movement of IS elements. The ability of *E. faecium* to rapidly diversify may contribute to its success as a nosocomial pathogen as it allows clones that circulate in a hospital to rapidly optimize their ability to effectively colonize individual patients that may differ in their underlying illnesses, antibiotic therapy and composition of the gut microbiota. Improving our understanding of the mechanisms that underpin this trait is crucial for combating the issues related to the emergence of multidrug-resistant *E. faecium* as an important nosocomial pathogen.

## Methods

### Strain collection

Ninety-six *E. faecium* strains were isolated from five patients during routine diagnostic screenings at the University Medical Center Utrecht, a tertiary care facility in Utrecht, the Netherlands, as part of routine screening for colonization by multidrug-resistant *E. faecium* [26, 42]. Patients were screened for carriage of multidrug-resistant *E. faecium* by culturing rectal swabs in Enterococcosel broth (Becton Dickinson) supplemented with aztreonam (75 mg/liter) at 37°C. If the cultures exhibited black colorization within 48 h, the broth was streaked on an Enterococcosel Agar plate (Becton Dickinson), supplemented with aztreonam and vancomycin (25 mg/liter) or with aztreonam and ampicillin (16 mg/liter) and incubated at 37°C for 48 h. Black colonies formed by Gram-positive cocci were subjected to multiplex PCR to detect vancomycin resistance genes and the *esp* gene, as well as additional antibiotic susceptibility testing. If a vancomycin- and/or an ampicillin-resistant isolate was found during a screening, this isolate was subsequently stored at −80°C. These patients were selected due to their relatively high number of available screening isolates (between 17 and 25 per patient). One patient (patient C) was admitted to the hospital for recurring abscesses on the upper leg, the other four patients were admitted for (hemo)dialysis procedures. None of the patients were diagnosed with enterococcal infections.

### Growth curves and maximum growth rate

A BioScreen C instrument (Oy Growth Curves AB) was used to measure bacterial growth. One colony was picked per strain and grown overnight in Brain Heart Infusion (BHI) broth at 37°C with shaking at 200 rpm, then diluted to an initial optoical density at 600 nm (OD_600_) of 0.1 in BHI. The cultures were incubated in triplicate in the Bioscreen C system at 37°C with continuous shaking, and absorbance at 600 nm (A_600_) was recorded every 15 min for 9 hours. The growth rates (*μ*) were calculated using 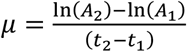, where t_x_ signifies a time point and A_x_ the associated A_600_ at this time point. The maximum growth rate (*μ_max_*) was determined for each individual experiment by taking the highest *μ* over the course of the growth.

### DNA isolation, genome sequencing and assembly

Genomic DNA of all strains was isolated from overnight cultures in Brain Heart Infusion broth, incubated at 37°C with shaking at 200 rpm, using the Wizard Genomic DNA purification kit (Promega). Library preparation for sequencing was done using the Nextera XT kit and 150 nucleotide paired-end sequencing was performed by Edinburgh Genomics on an Illumina HiSeq 2500. An additional 70 publicly available *E. faecium* genomes, described in [8], were also included in our analyses and were used to represent the global diversity of the species *E. faecium*. The Nesoni (version 0.122) tool [43] was used to remove adapter sequences and homopolymers, and to trim low-quality bases in sequence reads that had a quality score below 10. If more than half of a read was composed of low-quality bases, the read was discarded. The SPAdes assembler (version 3.1.0) [44] with *--careful* option and with k-mer sizes of 21, 31 and 41 was used for genome assembly. From the resulting contigs, those with less than 10-fold nucleotide coverage, as well as those smaller than 500 bases were discarded.

Assembly quality was checked using QUAST [45] and contigs not originating from bacteria (presumably due to low-level contamination of datasets with eukaryotic reads) were identified by aligning to NCBI GenBank database using BLASR+ (version 2.2.29) [46] and were removed.

The genome of strain A_020709_82 (GenBank accession number CP018128) was sequenced to serve as a reference for the analysis of the accessory genomes and distribution of IS elements in the genomes of the strains in group 1. DNA was prepared as described above, and then prepared for sequencing according to the Genomic DNA sequencing for the MinION device protocol (Oxford Nanopore Technologies, March 2016). From the obtained Pre-Sequencing Mix, approximately 60ng was loaded on a R7.3 flowcell and sequenced using an Oxford Nanopore MinION MkI instrument, which was run for a total of 48 hours with a Pre-Sequencing Mix top up (~60ng) at the 24-hour mark. A total of 18,629 high-quality two-directional (2D) reads were produced for a total of ~127 million bases. Poretools [47] was used to extract a fasta-format file containing the reads. A hybrid assembly using these reads combined with 2 x 150 bp HiSeq 2500 Illumina reads was then generated using SPAdes 3.7.0 [44] with the -- nanopore option.

### Genome annotation and clustering of orthologous proteins

We annotated the genome assemblies of all 166 isolates included in this study by using the Prokka [48] annotation tool (version 1.10) with its default parameters. To create clusters of orthologous proteins, the amino acid sequences of all genes in the 166 genomes were aligned against themselves using BLAST+ (version 2.2.29) [46]. Orthologous genes were identified with orthAgogue (version 1.0.3) [49] using the bit score information from the BLAST alignments, where aligned sequence length between two genes should at least be half of the size of the longer gene. Orthologous genes were grouped into OGs using the MCL algorithm (version 12-135) with the inflation parameter of 1.5 [50].

### Phylogenetic analyses

We generated core genomes by concatenating the sequences of OGs that were present once in all genomes. To prevent bias in our data created by recombination, we filtered the core genomes to identify putative recombination regions using the Gubbins recombination filtering tool (version 1.3.4) [51]. We then used the SNPs in the core genome located outside of the identified recombination regions to create two phylogenetic trees: one for all 166 strains (96 patient isolates and 70 publicly available genomes), and one for the 114 clade A1 strains (95 patient isolates and 19 clade A1 isolates as defined in [8]), using FastTree2 (version 2.1.7; double precision mode enabled) [52]. We used a GTR substitution model for nucleotide sequences with a Gamma site evolutionary rate correction and 1000 bootstrap samples to estimate the support for bifurcation points.

We observed 6 different groups of strains among newly sequenced 96 strains based on their similarity in the tree of 114 clinical isolates. Each of these 6 groups was then analyzed separately. For each group of strains, OG clustering as well as recombination filtering was applied as described above. However, instead of using SNPs, a concatenated core genome containing all the core genes that are outside of recombination regions were used to obtain more accurate branch lengths and better estimates of time divergence in phylodynamic analysis [53].

### Estimation of mutation rates

To further analyze the evolutionary dynamics of each group of strains, we first checked for the presence of sufficient temporal signal (R^2^>0.30) using Path-O-Gen (version 1.4pre) [28]. We then used the BEAST molecular evolutionary analysis tool (version 1.8.2) [54] only for those groups that had a sufficient level of temporal signal.

We used jModelTest2 [55] to identify the substitution model and site heterogeneity model, and to estimate the proportion of invariant sites, the transition/transversion ratio and the shape parameter of the Γ distribution. Five different clock models (strict, exponential, log-normal, fixed and random) and three different demographic models (constant, log-normal and Bayesian-skyline plot) were used in the BEAST analysis. These different models were analyzed with 100 million Markov chain Monte Carlo (MCMC) simulations with 10 million burn-ins, where sampling was done after every 10,000 simulations. The best model among these 15 models (5 clock models x 3 demographic models) was selected using path sampling (PS) and stepping-stone sampling (SS) model selection algorithms with one million simulations and 100 path steps, where logs after every 1000 simulations were screened as described previously [56]. Maximum clade credibility (MCC) tree was generated using TreeAnnotator using the median heights of trees [54]. The estimated prior values by jModelTest2 for substitution and heterogeneity models were HKY and I+G for both groups. The rest of the estimated coefficients were the same with the exception of the transition/transversion ratio being 6.79 and 3.88 for group 1 and 4, respectively. The best BEAST model for group 1 isolates was a lognormal relaxed clock (lognormal) with a constant coalescence model (lognormal-constant) based on SS model selection, and a lognormal relaxed clock with a Bayesian skyline (BS) coalescence model (lognormal-BS) based on the PS model selection. Although the SS model selection method is generally more accurate than the PS model selection method [57], we chose the lognormal-BS as the BS coalescence model had higher effective sample size values than the constant model; and the difference between lognormal-constant and lognormal-BS models was negligible. For group 4 strains, the best BEAST model was the lognormal relaxed clock with an exponential coalescence model according to both the SS and PS model selection methods.

### Analysis of accessory genome

Besides core-genome based analysis, we studied the differential presence of accessory genes within the six groups. When an OG was either present in less or absent in more than 90% of the strains in the group, it was included in the accessory genome. In addition to the annotation information, we also considered on which contigs differentially present genes were located in order to identify potential genetic links. Thus, we aligned the corresponding contigs of each differentially present gene against the GenBank database using BLAST+ (version 2.2.29+) [46] to identify putative mobile elements on which the variably present genes were located. We aligned differentially present genes of group 1 to the A_020709_82 reference genome and visualized location of these genes on the reference using Circos (version 0.69) [58]. The two largest clusters were aligned to all 166 genomes using BLAST+ (version 2.2.29+) [46] to determine the presence/absence of these regions in *E. faecium*. Abricate [27] was used to determine the presence of antimicrobial resistance determinants in the assembled genomes.

### Gain and loss of insertion sequences

We used ISMapper [29] to find gain and loss of insertion sequence elements (IS elements) among group 1 strains, which solely includes isolates from the same patient (patient A). Sequences of IS elements were found by uploading the complete genome sequence of the A_020709_82 reference strain at the ISfinder [59] website. Sequences of the identified IS elements were used in ISmapper together with the sequence reads from patient isolates. The genome of the patient A strain A_020709_82 was used as a reference to which reads were aligned, positions of IS elements were ordered regarding their positions in the reference genome.

## List of abbrevations

SNPs: single nucleotide polymorphisms
IS: insertion sequence
HGT: horizontal gene transfer
WGS: whole genome sequencing
ARE: ampicillin-resistant *E. faecium*
VRE: vancomycin-resistant *E. faecium*
HPD: highest posterior density
OGs: orthologous genes
BHI: Brain Heart Infusion
OD_600_: optical density at 600 nm (OD_600_)
A600: absorbance at 600 nm (A_600_)
µ: growth rate
μ_max_: maximum growth rate
MCMC: Markov chain Monte Carlo (MCMC)
PS: path sampling
SS: stepping-stone sampling
MCC: maximum clade credibility
BS: Bayesian skyline
ENA: European Nucleotide Archive
ICU: Intensive Care Unit

## Declarations

Ethics approval and consent to participate

Strains were isolated as part of routine diagnostic procedures during a VRE outbreak. This aspect of the study did not require consent or ethical approval by an institutional review board.

Consent for publication Not applicable

Availability of data and material

Short read data for the 96 genomes sequenced in this study are available at the European Nucleotide Archive (ENA), accession number PRJNA344739. The long-read sequence dataset used for the assembly of the genome of strain A_020709_82 is available at ENA, accession number CP018128.

**Competing interests**
The authors declare that they have no competing interests

## Authors’ contributions

W.v.S. designed the study. A.T. provided the strains. J.R.B., J.B., M.R.C.R. performed data analyses. J.R.B., J.B., R.J.L.W., and W.v.S. wrote the manuscript. All authors read and approved the final manuscript.

## Acknowledgements

The authors wish to thank Mick Watson (Edinburgh Genomics, University of Edinburgh) for generating the Illumina sequence data used in this study.

