## Supplemental Figures for "*Enterococcus faecium* genome dynamics during long-term asymptomatic patient gut colonization"

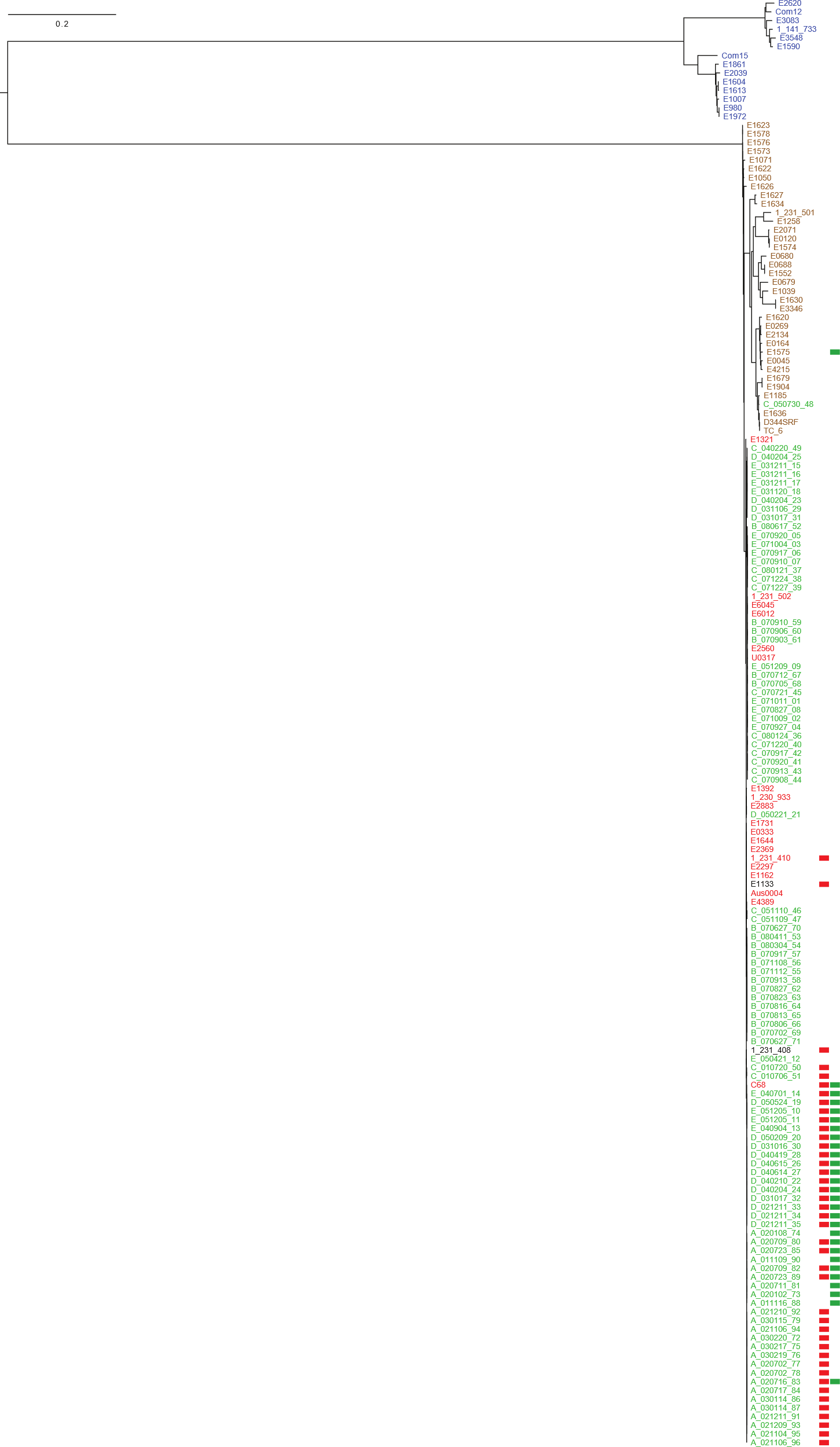

**Supplementary figure 1**. Core genome-based, recombination-filtered phylogeny of all 166 isolates in this study. Clade B strains are indicated in blue, clade A1 strains are indicated in red, clade A2 strains are indicated in brown. Strains that were sequenced as part of this study are indicated in green. Red and green blocks mark the presence of the phage element (cluster 1) and the carbohydrate metabolism-related element (cluster 2), respectively, which are discussed in the main text.

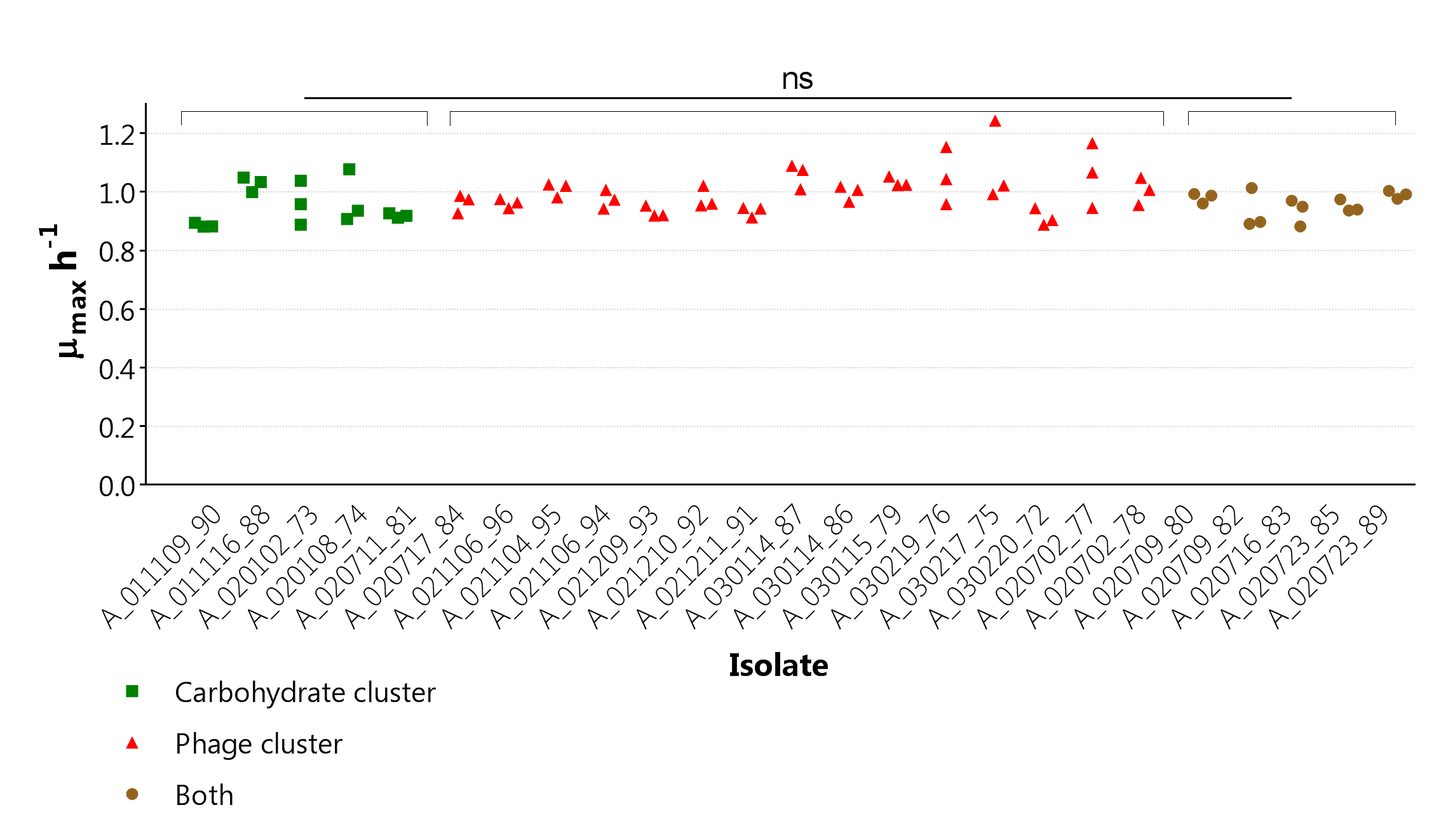

**Supplementary figure 2:** Maximum growth rates (*μ_max_*) of the 25 patient A isolates. Isolates were cultured in triplicate. The colours indicate which variably present genetic elements (further described in the text) the isolates contain: green, the carbohydrate metabolism-related element (green squares), the phage element (red triangles) or both elements (brown circles). No significant differences between groups were found
